# Capturing *in vivo* RNA transcriptional dynamics from the malaria parasite *P. falciparum*

**DOI:** 10.1101/099549

**Authors:** Heather J. Painter, Manuela Carrasquilla, Manuel Llinás

**Affiliations:** Department of Biochemistry & Molecular Biology and Center for Malaria Research, Pennsylvania State University, University Park, PA 16802; current address Wellcome Trust Sanger Institute, Wellcome Genome Campus, Hinxton, Cambridge, CB10 1SA, UK

## Abstract

To capture the transcriptional dynamics within proliferating cells, methods to differentiate nascent transcription from pre-existing mRNAs are desired. One approach is to label newly synthesized mRNA transcripts *in vivo* through the incorporation of modified pyrimidines. However, the human malaria parasite, *Plasmodium falciparum*, is incapable of pyrimidine salvage for mRNA biogenesis. To capture cellular mRNA dynamics during *Plasmodium* development, we have engineered parasites that can salvage pyrimidines through the expression of a single bifunctional yeast fusion gene, cytosine deaminase/uracil phosphoribosyltransferase (FCU). We show that expression of FCU allows for the direct incorporation of thiol-modified pyrimidines into nascent mRNAs. Using developmental stage-specific promoters to express FCU-GFP enables the biosynthetic capture and in-depth analysis of mRNA dynamics from subpopulations of cells undergoing differentiation. We demonstrate the utility of this method by examining the transcriptional dynamics of the sexual gametocyte stage transition, a process that is essential to malaria transmission between hosts. We find that sexual stage commitment is governed by transcriptional reprogramming and the stabilization of a subset of essential gametocyte transcripts. This new method for biosynthetic labeling of *Plasmodium* mRNAs is incredibly versatile and can be used to measure transcriptional dynamics at any stage of parasite development, and thiol-modified RNAs will allow for future applications to measure RNA-protein interactions in the malaria parasite.

## INTRODUCTION

*Plasmodium falciparum*, the causative agent of malaria, has a complex life cycle that includes development in multiple tissues within the human host and the female *Anopheles* mosquito vector (Bannister and Mitchell 2003). A mosquito initiates infection by injecting *Plasmodium* sporozoites into a human, which migrate to the liver where they undergo asexual replication (schizogony) forming thousands of merozoites. When merozoites are released from the liver they initiate the blood stage of infection. The intraerythrocytic development cycle (IDC) consists of continuous 48-hour cycles of maturation and cell division, during which up to 32 new daughter cells are formed and released each cycle. These clonal progeny invade uninfected red blood cells and repeat the asexual replication process, exponentially increasing the population size. During the IDC a small proportion of asexual parasites transition to the sexual stage and differentiate into female or male gametocytes (Guttery et al. 2015). Gametocytogenesis is a stochastic process that is obligatory for parasite transmission, as asexual forms cannot propagate within the mosquito vector (Josling and Llinas 2015). These gamete precursors are morphologically and functionally distinct from their asexual blood stage counterparts, which is reflected in their cellular development, metabolism, and patterns of gene expression (Liu et al. 2011).

Gene expression in eukaryotic systems is comprised of many levels of regulation including chromatin modification, active transcription mediated by *trans*-acting factors, and post-transcriptional mRNA turnover or stabilization (Lelli et al. 2012). Similarly, *Plasmodium* differentiation and growth within various cell types and hosts involves complex regulatory mechanisms that govern both transcriptional and post-transcriptional processes (Hughes et al. 2010; Cui et al. 2015; Vembar et al. 2016). Development during the IDC is driven by a coordinated transcriptional cascade with the majority of genes expressed in a “just-in-time” fashion (Bozdech et al. 2003; Le Roch et al. 2003), although the mechanisms that underlie the coordination and specificity of active transcription remain largely uncharacterized (Hughes et al. 2010; Painter et al. 2011). Thus far, only a single transcription factor family, the 27-member Apicomplexan AP2 (ApiAP2) proteins, has emerged as transcriptional regulators with functions across all developmental stages (Balaji et al. 2005; Painter et al. 2011; Iwanaga et al. 2012; Josling and Llinas 2015; Yuda et al. 2015). Given the task of regulating roughly 5,500 *Plasmodium* genes with such a small repertoire of transcription factors, it has been proposed that mRNA dynamics in *Plasmodium* parasites are greatly influenced by post-transcriptional regulatory mechanisms (Hughes et al. 2010; Bunnik et al. 2016; Vembar et al. 2016).

Evidence for post-transcriptional regulation in *Plasmodium* spp. includes several studies that have reported a significant delay between transcription and translation (Le Roch et al. 2004; Hall et al. 2005; Foth et al. 2008; Foth et al. 2011), and recent studies have demonstrated ribosomal influence on the timing of mRNA translation (Bunnik et al. 2013; Caro et al. 2014). Despite these insights, the proteins that regulate these processes are largely unknown. Surprisingly, although bioinformatic analyses have suggested that between 4-10% of the *Plasmodium* genome encodes RNA-binding proteins (RBPs), only a handful have been characterized to date (Reddy et al. 2015; Bunnik et al. 2016). Evolutionarily conserved post-transcriptional regulatory factors such as DOZI (DDX-6 class DEAD box RNA helicase), CITH (Sm-like factor homolog of CAR-I and Trailer Hitch) and a pumilio family protein (PUF2) play critical roles in the translational repression (TR) of essential genes throughout *Plasmodium* sexual development (Cui et al. 2015; Vembar et al. 2016). Other important RBPs are the Alba-domain containing proteins (PfAlba1-4) which interact with RNA throughout the parasite’s lifecycle and are found associated with TR-complexes (Chene et al. 2012; Vembar et al. 2015). Although the role of *Plasmodium* RBPs in gene regulation is a growing area of interest, techniques to accurately measure mRNA dynamics and identify post-transcriptional regulatory factors in *Plasmodium* parasites lag far behind those available for other eukaryotic systems.

In recent years, the ability to incorporate thiol-modified pyrimidines into nascent mRNA transcripts from human cells (Cleary et al. 2005), mice (Kenzelmann et al. 2007; Gay et al. 2013), *Drosophila* (Miller et al. 2009), and yeast (Miller et al. 2011; Munchel et al. 2011; Neymotin et al. 2014) has afforded an effective tool to evaluate mRNA dynamics of diverse cellular populations, both spatially (Miller et al. 2009; Gay et al. 2013) and temporally (Miller et al. 2011; Munchel et al. 2011). However, thiol-modified mRNA capture is dependent upon pyrimidine salvage, and the evolutionary lineage of *Plasmodium* has lost this biochemical capacity (Hyde 2007). Genetic supplementation of pyrimidine salvage enzymes in *Plasmodium* would restore this metabolic pathway, thereby enabling methods that utilize biosynthetically labeled RNA (Friedel and Dolken 2009)

To probe transcriptional dynamics in *P. falciparum*, we have developed a customizable approach to capture active transcription and mRNA stabilization in the parasite. Our method is made possible by the exogenous expression of two enzymes involved in pyrimidine salvage: the *Saccharomyces cerevisiae* gene <u>f</u>usion of <u>C</u>ytosine Deaminase and <u>U</u>racil Phosphoribosyltransferase (FCU). With an active pyrimidine salvage pathway, *P. falciparum* can readily uptake biosynthetically modified pyrimidines and incorporate these into nascent transcripts, which can be tracked at various time points or in specific cells throughout parasite development. These transcripts can subsequently be captured and analyzed with common comprehensive transcriptomic approaches such as DNA microarrays or RNA-seq. This mRNA capture method from FCU-expressing parasites can be used to temporally profile the transcriptional dynamics of the developing *Plasmodium* parasite at all stages of development.

As a proof of concept, we used biosynthetic mRNA capture to measure mRNA dynamics from the small fraction of *Plasmodium* parasites undergoing commitment to gametocytogenesis. Capturing the transcriptional program of early gametocytes is difficult because of technical challenges associated with isolating and distinguishing early stage gametocytes from asexual trophozoites (Sinden 1982; Silvestrini et al. 2010). To circumvent these challenges, we generated parasites lines engineered to express FCU-GFP using the established gametocyte-specific *pfs16* (PF3D7_1302100/PFD0310w) promoter, which is only actively transcribed and expressed in a subset of parasites committed to gametocytogenesis (Bruce et al. 1994; Dechering et al. 1997; Dechering et al. 1999) and which restricts the biosynthetic capture of 4-thiouracil modified mRNA to early gametocytes. Our methodology does not require physical isolation of the early gametocyte subpopulation, even though it identifies the sexual-stage mRNA dynamics associated with this developmental differentiation process. Using this approach, we find that regulation of early sexual stage development is dictated by a combination of transcriptional reprogramming and enhanced stabilization of gametocyte stage-specific transcripts.

## RESULTS

### Addition of a functional pyrimidine salvage pathway into *Plasmodium*

The measurement of transcriptional dynamics is greatly facilitated by the exogenous incorporation of modified pyrimidines, which allows for the biosynthetic labeling, capture, and subsequent analysis of newly transcribed RNAs (Fig. 1A). *Plasmodium* parasites are naturally unable to incorporate pyrimidine nucleoside precursors via salvage and instead rely on *de novo* pyrimidine biosynthesis (Reyes et al. 1982). To enable parasites to salvage pyrimidines, we genetically modified the *P.f.* 3D7 laboratory strain to exogenously express a functional *S. cerevisiae* fusion gene of cytosine deaminase (*FCY1*) and uracil phosphoribosyltransferase (*FUR1*) (Erbs et al. 2000), with a C-terminal GFP-tag (FCU-GFP) under the control of a constitutively active *P.f. calmodulin* promoter (*cam*, PF3D7_1434200) (Crabb and Cowman 1996) (Fig. 1B). Transgenic 3D7^*cam*^ parasites, but not wild-type parasites, readily incorporated the thiolated pyrimidine precursor 4-TU in a concentration dependent manner (3D7) (Fig. 1C).

**Figure 1:**
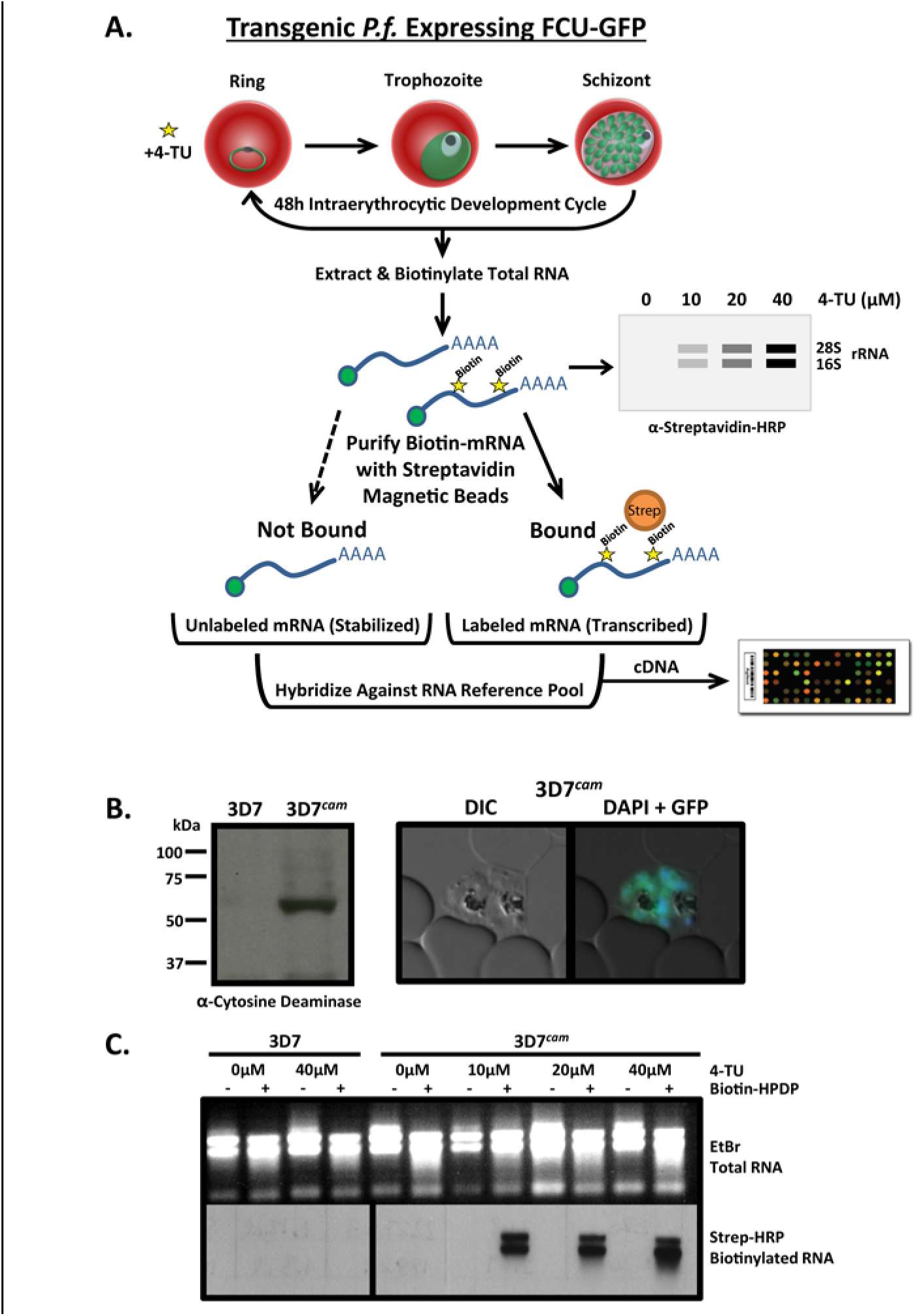
Engineering *P. falciparum* to salvage pyrimidines and generate thiol-modified RNAs. A) Schematic of 4-TU biosynthetic mRNA capture method. Transgenic *P. falciparum* expressing a fusion gene containing cytosine deaminase/uracil phosphoribosyltransferase tagged with GFP (FCU-GFP) under the control of the calmodulin (CAM) promoter (3D7^*cam*^) enables 4-TU salvage and incorporation into RNA. Thiolated-RNA can be biotinylated and detected by Northern blot or affinity purified by streptavidin magnetic beads for analysis by DNA microarray. B) Expression of FCU-GFP from 3D7^*cam*^ was verified by Western blot when probed with anti-yeast cytosine deaminase and by live fluorescence microscopy (GFP = green, nuclear DNA stained with DAPI = blue) C) Both wild-type and 3D7^*cam*^ parasites were grown for 12 hrs in the presence of increasing 4-TU concentrations. The specificity of RNA thiol-incorporation and biotinylation was assessed by running 2 μg of each RNA sample with and without EZ-link Biotin-HPDP incubation (top panel). Total RNA was transferred to a nylon membrane and probed with streptavidin-HRP to detect biotinylated RNAs (bottom panel).

To ensure that cyclical progression through the 48-hour IDC was not detrimentally affected by the incorporation of thiol-modified pyrimidines, 3D7^*cam*^ parasites were incubated in the presence of increasing concentrations of 4-TU (0, 20, 40, 80, 160μM), while monitoring parasitemia for 72 hours (Fig. S1A). Based on these data, we selected 40μM 4-TU for further experimentation, since concentrations at or below this value had no effect on parasite growth (Fig. S1A). Global transcriptional profiling by DNA microarray analysis of a complete 48-hour cycle of 3D7^*cam*^ grown in 40μM 4-TU further indicated that developmental progression is unperturbed (Fig. S1B). To ensure that the uptake and incorporation of 4-TU was consistent throughout development, pulses of 4-TU (40μM) were administered at the ring (10hr), trophozoite (24hr), and schizont stages (38hr) for varying lengths of time (0, 1, 2, and 4hrs) (Fig. S1C). Our results indicated that 3D7^*cam*^ parasites were fully capable of transporting and incorporating 4-TU into the total RNA pool at all stages. Nascent transcripts that incorporated 4-TU were readily isolated from total RNA through covalent biotinylation of the thiol-labeled RNAs, followed by affinity purification (Fig. 1A and S1D).

To ensure robust incorporation of 4-TU into the pyrimidine pool, we determined the efficiency of the transgenic pyrimidine salvage pathway versus endogenous *de novo* pyrimidine synthesis (Fig. 2A). Using liquid chromatography-mass spectrometry (LC-MS), we directly measured the incorporation of ^15^N_2_-uracil into the uracil monophosphate (UMP) pyrimidine pool. The results clearly show that 3D7^*cam*^ parasites expressing FCU-GFP incorporated heavy-labeled uracil (^15^N2-UMP) into 50% of the pyrimidine pool in 10 minutes (Fig. 2A and 2B). We next tested whether 3D7^*cam*^ parasites could grow in the absence of *de novo* biosynthesis, by using the antimalarial drug atovaquone to inhibit the cytochrome *bc_1_* complex in the mitochondrial electron transport chain (mtETC). Atovaquone treatment blocks ubiquinone regeneration, which in turn leads to an inhibition of dihydroorotate dehydrogenase (DHOD), an essential enzyme in pyrimidine *de novo* biosynthesis requiring ubiquinone (Fig. 2A) (Painter et al. 2007). 3D7^*cam*^ parasites were grown in the presence of atovaquone (10xIC_50_) for 3 hours, followed by supplementation of the medium with ^15^N_2_-uracil and ^13^C-bicarbonate. We found that these parasites readily salvaged the exogenously supplied pyrimidine (^15^N_2_-UMP), while *de novo* pyrimidine synthesis was disrupted by atovaquone (^13^C-UMP) (Fig. 2A and 2C). As expected, 3D7^*cam*^ parasites were able to bypass the effect of atovaquone when uracil was supplied as a pyrimidine precursor to the growth medium (Fig. 2D). Therefore, the introduction of the FCU enzyme resulted in parasites that rapidly and efficiently salvage pyrimidine precursors and can proliferate independent of *de novo* pyrimidine biosynthesis.

**Figure 2:**
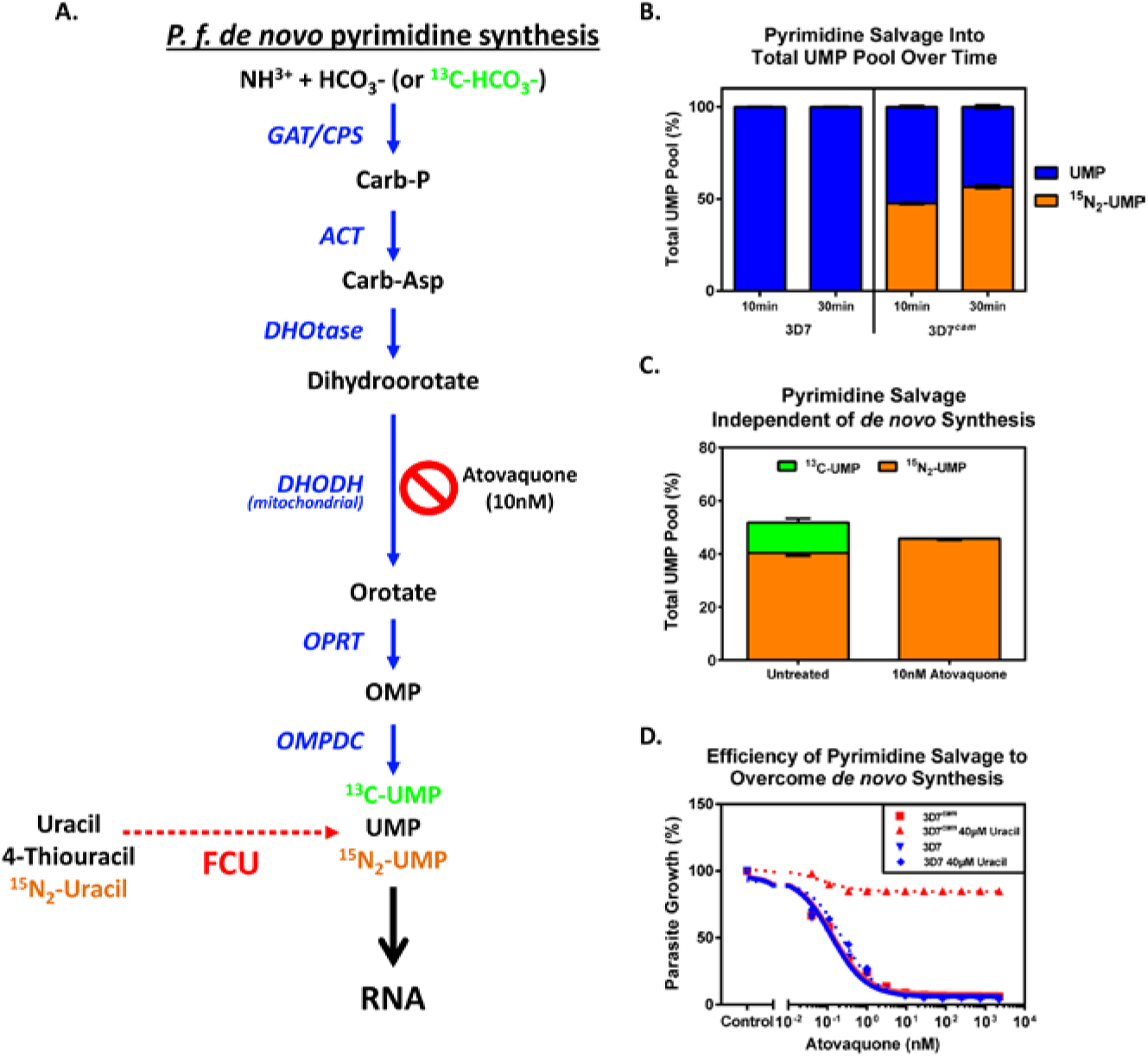
Efficiency of pyrimidine salvage in FCU-GFP expressing *P. falciparum*. A) In *Plasmodium* parasites, pyrimidines are metabolized *de novo* from bicarbonate and ammonia through a series of enzymatic reactions (blue arrows) resulting in the generation of UMP, which can be incorporated into nascent RNA (black arrow). FCU-GFP allows for salvage of the pyrimidine precursor uracil (red dashed arrow) to UMP, which can then be incorporated into RNA. Inhibition of *de novo* pyrimidine synthesis is achieved via treatment with the mitochondrial inhibitor atovaquone. B) 4-TU is efficiently taken up and incorporated into the pyrimidine pool by *P.f.* 3D7^*cam*^ parasites grown in the presence of 40μM ^15^N_2_-uracil for 10 and 30min. Cellular metabolites were detected via LC-MS and the proportion of total UMP pool that is unlabeled (blue) and labeled with ^15^N_2_ (orange) was calculated (n = 2 ± s.d. performed in triplicate). C) Flux through *de novo* pyrimidine synthesis and salvage was measured in treated and untreated parasites by LC-MS after the addition of ^13^C-bicarbonate (^13^C-HCO_3_-) and ^15^N_2_-uracil to the culture medium for 30 min. The percentage of UMP derived from *de novo* synthesis (^13^C-UMP) and salvage (^15^N_2_-UMP) was determined in the presence or absence of the inhibitor atovaquone (n = 2 ± s.d. performed in triplicate). D) Early ring stage *P. falciparum* 3D7 and 3D7^*cam*^ were exposed to titrating concentrations of atovaquone with or without supplementation of 40 μM uracil to the medium. Parasite survival was determined using a traditional 48 hr SYBR-green growth inhibition assay and plotted as an average of three technical replicates (± s.d.).

### Stage-specific expression of pyrimidine salvage allows for biosynthetic labeling of parasites committed to sexual development

The 3D7^*cam*^ parasite line ubiquitously expresses functional pyrimidine salvage enzymes in all cells from a relatively strong constitutive promoter. To demonstrate the utility and universality of the FCU-GFP mediated biosynthetic mRNA capture methodology in *P. falciparum*, we wanted to characterize the mRNA dynamics of a subpopulation of cells undergoing a unique developmental state. As a proof of concept, we focused on parasite differentiation from the asexual blood stage to the sexual stage (gametocyte). To enable biosynthetic mRNA capture during the developmental transition to sexual differentiation, we placed *fcu-gfp* under the control of the *pfs16* promoter (3D7^*pfs16*^) (Dechering et al. 1999; Pradel 2007; Eksi et al. 2008; Adjalley et al. 2011) (Fig. 3A, 3B and S2A). By regulating expression using *pfs16*, the FCU-GFP fusion protein should be expressed only in the subpopulation of cells that are committed to become gametocytes, thereby enabling 4-TU salvage exclusively from these cells (Fig. 3A). As anticipated, 3D7^*pfs16*^ parasites have a reduced number of GFP positive cells compared to the 3D7^*cam*^ parasite population (Fig. 3A and S2B). Direct quantification of 4-TU incorporation into cellular RNA from 3D7^*cam*^ vs. 3D7^*pfs16*^ parasites also demonstrated a significant reduction in 3D7^*pfs16*^ (Fig. 3C and S2C). As observed in the 3D7^*cam*^ parasites, mRNA abundance profiles of 3D7^*pfs16*^ throughout the IDC were unperturbed when grown in the presence of 4-TU (Fig. S1B). Therefore, we conclude that *pfs16*-FCU-GFP is expressed only in a sub-population of parasites undergoing sexual differentiation (Fig. 3A).

**Figure 3:**
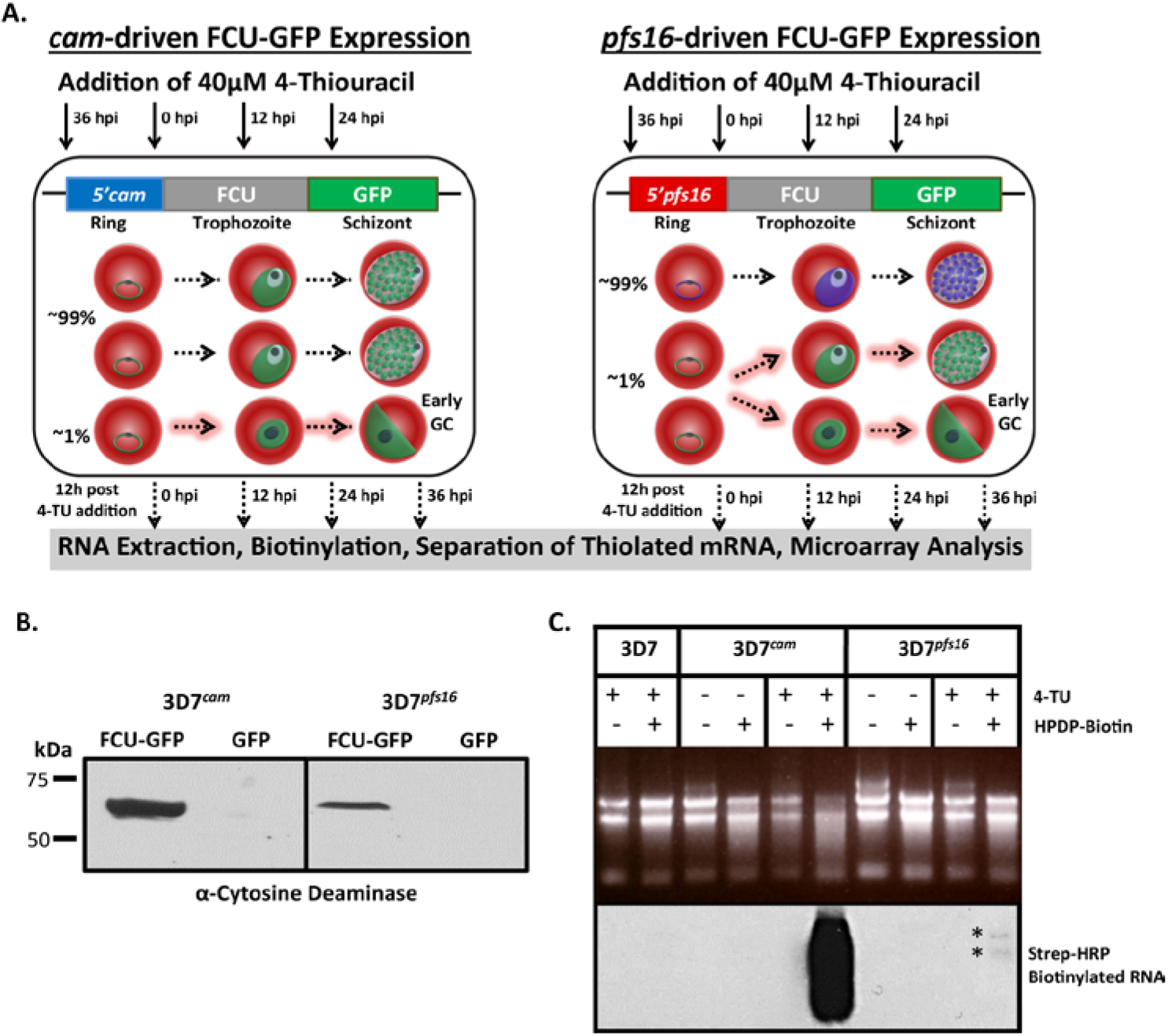
Stage-specific pyrimidine salvage for detection of early gametocyte transcription. To measure mRNA from a subpopulation of cells, we expressed FCU-GFP under the control of the gametocyte-specific promoter *pfs16* (3D7^*pfs16*^). A) Schematic representation of the experimental design including the timing of 4-TU incubation (black arrows) and RNA extraction (dashed black arrows), plasmids transfected into *P. falciparum* strains, and a depiction of the highly synchronous cell populations that express *cam*- and *pfs16*-FCU-GFP throughout the 48 h IDC. All parasites express the constitutive *cam*-FCU-GFP (green) regardless of developmental stage (left panel, 100% GFP+). Only a small proportion (~1% GFP+) of parasites committed to sexual development express *pfs16*-FCU-GFP (green) while asexual parasites do not (right panel, ~99% GFP-). B) FCU-GFP protein from uninduced asexual cultures of 3D7^*cam*^ and 3D7^*pfs16*^ measured by Western blot probed with anti-yeast cytosine deaminase. C) Detection of subpopulation FCU-mediated thiol-tagged RNA was carried out by incubating highly synchronized *P.f.* 3D7, 3D7^*cam*^ and 3D7^*pfs16*^ in the presence or absence (top panel) of 4-TU for 12 hrs. Total RNA was extracted, biotinylated (top panel), and assayed by Northern blot probed with Streptavidin-HRP (bottom panel) demonstrating that thiol-tagging occurs at a much lower level in 3D7^*pfs16*^ (*) than in 3D7^*cam*^, representing the minor sexual-stage parasite subpopulation.

To capture the transcriptional dynamics associated with early gametocyte formation in *P. falciparum*, we compared the gametocyte producing 3D7 parasite line to the non-gametocyte producing F12 clone of 3D7 (Alano et al. 1995). The F12 parasite line is developmentally blocked due a loss-of-function nonsense mutation in the *pfap2-g* transcriptional regulator that prevents mature gametocyte formation (Kafsack et al. 2014) (Fig. S2D). Therefore, any significant differences detected in the transcriptional dynamics of 3D7 and F12 would be attributable to gametocyte commitment and development. To do this, the F12 parasite strain was modified to express FCU-GFP driven by either the *pfs16* or *cam* promoters (Fig. 3A). Expression of FCU-GFP within each strain was verified and all strains actively incorporate pyrimidines through salvage (Fig. 3B, 3C, S2A, S2B, S2C, S2E and S2F) enabling the temporal detection of differences in the mRNA dynamics during commitment to gametocytogenesis for both strains. Interestingly, the activity of the *pfs16*-promoter is independent of a parasite’s ability to form mature gametocytes (Fig. S2D), as F12^*pfs16*^ incorporated 4-TU through salvage albeit at significantly reduced levels compared to 3D7^*pfs16*^ (Fig. S2F).

### Stage-specific biosynthetic capture in various strains reveals asexual vs. gametocyte-specific mRNA dynamics

To measure the mRNA dynamics in early gametocytes from the 3D7 and F12 strains, we performed comparative biosynthetic mRNA capture (Fig. 1A) from synchronized *cam-* or *pfs16*-FCU-GFP expressing parasites at four time points throughout the 48-hour IDC (Fig. 3A) beginning with the ring stage of development. These highly synchronous parasites were separated into 4 equal fractions which were incubated with 40 μM 4-TU for a duration of 12 hours beginning at 36, 0, 12, and 24 hours post-invasion (hpi) (Fig. 3A). The first 12 hour incubation was initiated at 36 hpi, the point at which differentiation of asexual schizogony and stage I gametocyte development occurs (Bruce et al. 1990; Eksi et al. 2012) with the last extending to 36 hpi of the subsequent IDC. To enable the head-to-head comparison of the *cam*- versus *pfs16*-FCU-GFP expressing parasites, twelve hour incubations were used to ensure sufficient 4-TU incorporation into the small population of parasites that are committing to gametocytogenesis (*pfs16*-FCU-GFP positive, Fig. 3C and Fig. S2F). Following each 12 hour exposure, total RNA was extracted and biotinylated to separate thiol-labeled mRNAs (nascent transcription) from the unlabeled (stabilized) mRNAs via streptavidin-magnetic beads (Fig. 1A and 3A).

To identify differences in the mRNA dynamics of parasites committed to gametocytogenesis versus asexually replicating parasites, purified streptavidin-bound and unbound mRNAs from the four strains were analyzed using DNA microarrays at four timepoints (Fig. 3A). Microarray data from streptavidin-bound nascent mRNAs are referred to as “transcription” (containing newly incorporated 4-TU) (Fig. 4B) and the remaining unbound mRNAs (existing before 4-TU labeling) are referred to as “stabilization” (Fig. 4C). All genes identified by DNA microarray analysis were K10 means clustered based on the 3D7^*cam*^ and 3D7^*pfs16*^ strains (Fig. 4B and 4C, Table S1). An examination of the nascently transcribed genes identified from the *cam*-FCU-GFP expressing parasites demonstrates developmentally regulated gene expression of the *P. falciparum* transcriptome (Fig. 4B) similar to that previously reported from total mRNA abundance measurements (Fig. 4C) (Bozdech et al. 2003; Le Roch et al. 2003; Llinas et al. 2006). On the other hand, stabilized transcript dynamics follow a different periodic stage-specific pattern, demonstrating that nascent mRNA transcription and stabilization are independently regulated (Fig. 4C). Because the *cam* promoter is ubiquitously expressed in both asexual and sexual stage parasites, mRNAs from *cam*-FCU-GFP expressing parasites represent genes that are actively transcribed or stabilized in all parasites tested (Fig. 4A and S3A). Conversely, mRNAs from *pfs16*-FCU-GFP expressing parasites only represent the transcriptional dynamics of the subpopulation that are committing to gametocytogenesis (Fig. 4A and S3A).

**Figure 4:**
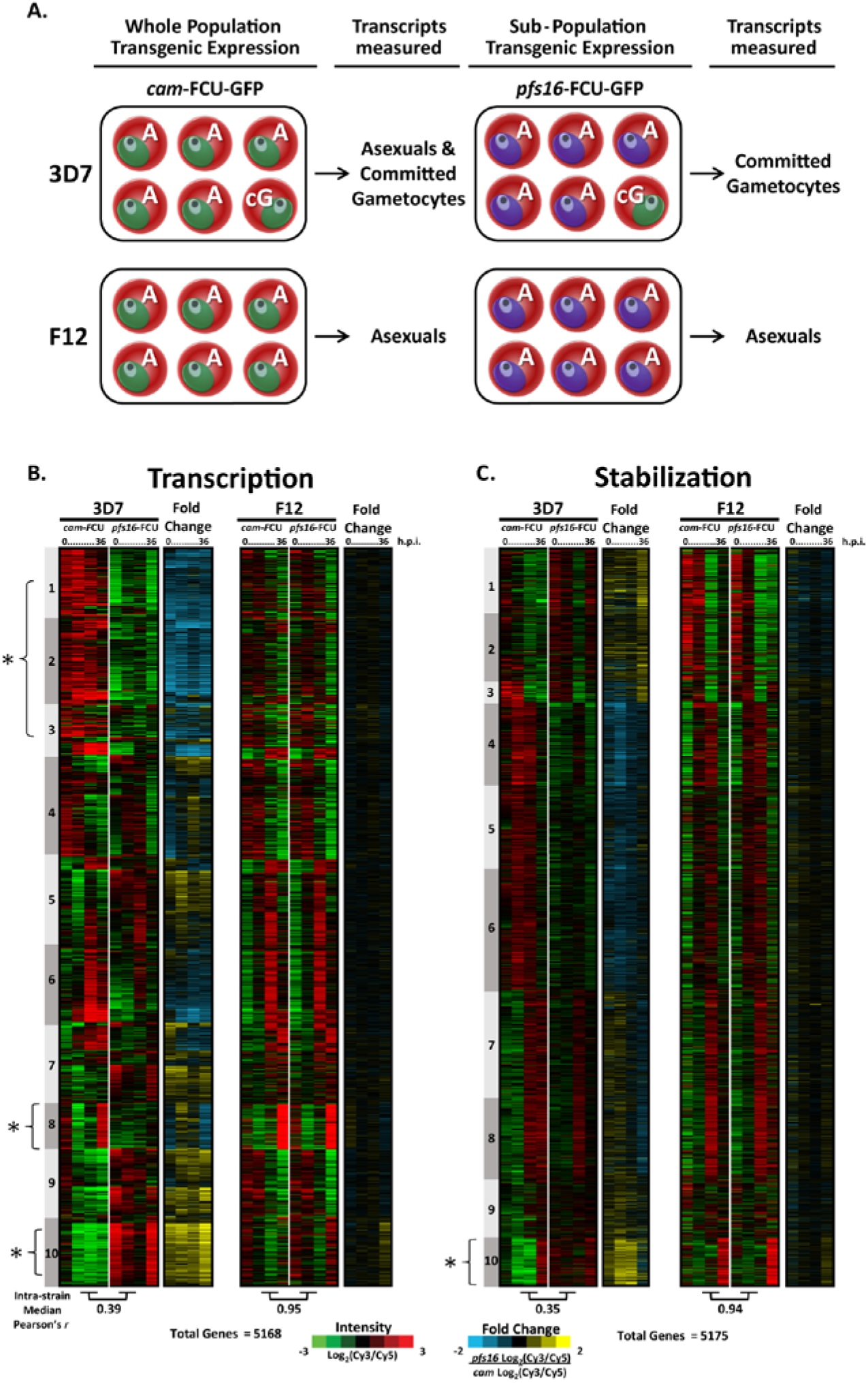
Biosynthetic mRNA capture of whole-genome thiol-labeled and non-labeled mRNAs during asexual and sexual development. A) Expression of *cam*-FCU-GFP occurs in the parasite strains 3D7 and F12 regardless of developmental stage. 3D7 produces gametocytes and the transcriptional dynamics in 3D7^*cam*^ are representative of both asexual (A) and committed gametocytes (cG). Expression of *pfs16*-FCU-GFP in 3D7^*pfs16*^ captures mRNA dynamics only in parasites committed to gametocytogenesis (cG). F12 is not able to produce mature gametocytes and mRNA dynamics occurring prior to gametocytogenesis can be measured regardless of the promoter used. B) Thiol-tagged RNA was separated from the total pool by streptavidin magnetic purification. mRNAs eluted from the beads are 4-TU labeled (representing “Transcription”) while C) unbound mRNAs were present before the addition of 4-TU (representing “Stabilization”). Each column represents the Log_2_(Cy3/Cy5) ratio for every gene in the sample detected above background for mRNAs transcribed (5168 genes) or stabilized (5175 genes) at 0, 12, 24, or 36 hours post invasion (h.p.i.). Ratios for each gene were K10 means clustered (1-10) and ordered according to their peak value starting at 0 h.p.i. The median Pearson’s *r* for each gene over time between strains is shown below the respective time-course. The fold change in transcription and stabilization for each gene (*cam*-FCU Log_2_(Cy3/Cy5)/*pfs16*-FCU Log_2_(Cy3/Cy5) was calculated and represented to aid in determining enrichment in the gametocyte-specific population. Clusters highlighted in the text are noted with an asterix.

We calculated the medium Pearson’s correlation at each of the IDC time-points for *cam*-FCU-GFP (asexual cells plus committed gametocytes) versus *pfs16*-FCU-GFP (committed gametocytes only) for both 3D7 and F12 parasites (Bottom, Fig. 4B, 4C,. S3C and S3D). These correlations directly reflect the strain’s ability to produce gametocytes (Fig. S2D). F12, which cannot produce gametocytes, has an almost perfect correlation for both newly transcribed and stabilized mRNAs, irrespective of the activity of either FCU-GFP promoter (transcription median *r* = 0.95, stabilization median *r* = 0.94) (Fig. S3B). In contrast, 3D7 has a significantly reduced intra-strain correlation (transcription median *r* = 0.39, stabilization median *r* = 0.35) (Fig. S3B). From these data, we concluded that the differences between *pfs16*-FCU-GFP and *cam*-FCU-GFP 3D7 parasite lines reveal a significantly altered gene regulatory program that reflects the parasite’s ability to commit to gametocytogenesis.

Upon closer examination of the 3D7 parasite strain data, we find several striking examples that reflect the difference between transcription in *pfs16*- and *cam*-FCU-GFP parasites. For example, Clusters 1 and 2 represent a large number of genes which are significantly down-regulated (or no longer transcribed) in *pfs16*-FCU-GFP 3D7 parasites. Of particular interest, is Cluster 8 (Fig. 4B, Table S1) which contains 320 genes that are highly transcribed in the *cam*-FCU-GFP strains and includes genes involved in merozoite egress and invasion (enrichment of GO:004409, entry into host, *p-value* = 9.69e-10) (Cowman et al. 2012) (Fig. S4, Table S1). These same transcripts are largely undetectable in the *pfs16*-FCU labeled pool (Fig. 4B, Cluster 8). For example, the gene encoding for merozoite surface protein 1 (*msp1*, PF3D7_0930300/PFI1475w) which is essential for asexual parasite invasion of erythrocytes, is transcribed at 36 hours in both F12^*cam*^ and F12^*pfs16*^ (Fig. S5A). However, *msp1* is only transcribed in 3D7^*cam*^ (Fig. S5A), presumably because sexually committed 3D7^*pfs16*^ parasites no longer require robust nascent transcription of this gene for reinvasion of erythrocytes. This suggests that we are measuring the transcriptional dynamics of a distinct subpopulation of parasites that no longer require active transcription of invasion ligands as might be expected for gametocytes, which develop for 10-12 days within the same red blood cell. Conversely, Cluster 10 (Fig. 4B, Table S1) contains 492 genes that are highly transcribed in 3D7^*pfs16*^, many of which have been previously reported as gametocyte genes (see below). For example, the gametocyte-specific gene *pf11-1* (PF3D7_1038400/PF10_0374) (Scherf et al. 1992) is highly transcribed in 3D7^*pfs16*^ at 36h, but not in F12^*pfs16*^ (Fig. S5A). Together, these data support the notion that we are capturing nascent transcription from the sexual stage subpopulation of parasites.

### Biosynthetic mRNA capture detects gametocyte-specific transcription and stabilization during sexual commitment

One of the goals of using our biosynthetic *pfs16*-FCU-GFP-based capture method was to detect genes transcribed early in gametocytogenesis. To identify these genes we calculated the fold change in the transcribed or stabilized signal intensities between the *pfs16*-FCU-GFP and *cam*-FCU-GFP expressing 3D7 and F12 parasite strains (Final column, Fig. 4B and 4C). The fold changes for the 3D7 parasite microarray data clearly show variation in the patterns of mRNA dynamics for virtually all transcripts measured (Fig. 4B and 4C). These ratios reveal that a large number of genes are no longer expressed in 3D7 gametocytes (Fig. 4B, ex: Transcription Clusters 1- 3, and 8), while other gene sets are strongly transcribed and stabilized (Fig. 4B and 4C, ex: Cluster 10). Using these data we defined a subset of 808 genes whose fold change ratio is higher in either the transcribed or stabilized data between 3D7^*pfs16*^ and 3D7^*cam*^ (Log_2_ fold change >1, 95^th^ percentile; Fig. 5A, Table S2). We hypothesized that these genes are likely to be involved in gametocyte commitment and development. To predict new genes associated with gametocytogenesis, we compared these gametocyte-enriched genes with previously published gametocyte datasets and found that many (672 of 808) of the genes enriched in *pfs16*-FCU-GFP parasites have been previously identified in gametocytes by RNA-seq (Stage II), DNA microarray (Stage II-IV), or proteomic (Stage I-V) analyses (Fig 5A) (Eksi et al. 2005; Silvestrini et al. 2005; Young et al. 2005; Mair et al. 2010; Silvestrini et al. 2010; Lopez-Barragan et al. 2011; Eksi et al. 2012; Brancucci et al. 2014; Tao et al. 2014). However, we found 136 genes that, to our knowledge, have not been associated with the gametocyte stage previously. Interestingly, among these genes, we found 35 rRNAs, 24 tRNAs, and 16 non-coding RNAs suggesting that the regulation of small RNAs may also play a role in the sexual differentiation of *Plasmodium* parasites (Table S2). While the majority of differences between 3D7^*cam*^ and 3D7^*pfs16*^ are reflected by an increase in transcription (Fig. 5A, Clusters 2-4), a smaller subset are stabilized (Fig. 5A, Cluster 1 and 5). These clusters of stabilized transcripts include a number of well-established gametocyte markers, but surprisingly also contains invasion genes such as *msp2*, *rhoph3*, and merozoite TRAP-like protein (*mtrap*) (Baum et al. 2006; Cowman et al. 2012) (Fig. 5A and Table S2). This finding support recent studies that have identified an essential role for MTRAP in mature gametocytes (Bargieri et al. 2016; Kehrer et al. 2016). Our results demonstrate the specificity and robustness of mRNA biosynthetic capture for measuring the mRNA dynamics from the parasite subpopulation undergoing gametocytogenesis.

**Figure 5:**
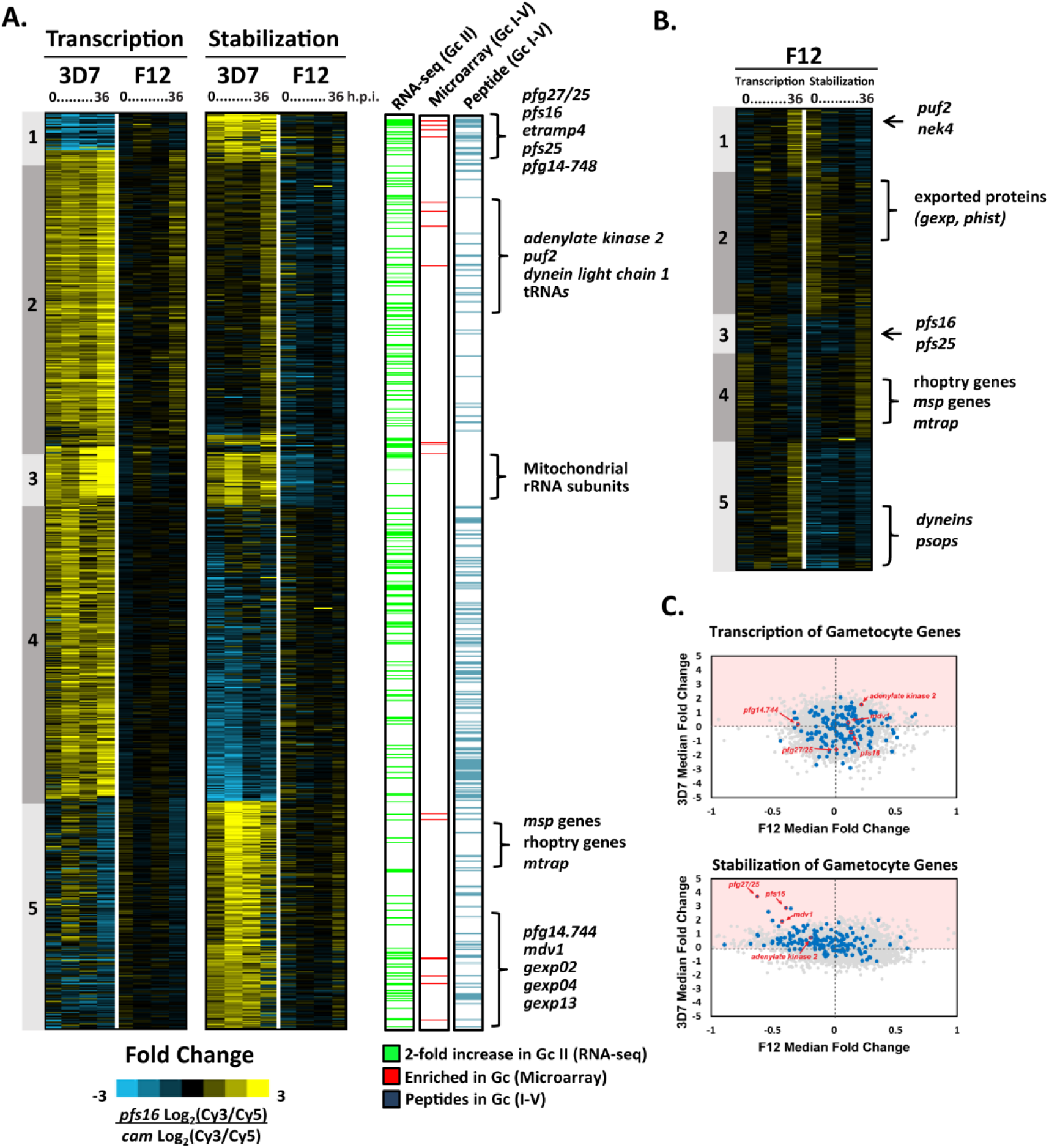
Dynamics of putative gametocyte-specific genes. A) Genome-wide gametocyte transcription and stability was calculated by determining the fold change in log_2_(Cy3/Cy5) ratio of *pfs16*-FCU-GFP/*cam*-FCU-GFP. A total of 809 genes enriched in *pfs16*-FCU-GFP expressing lines (≥ 1 log_2_ fold change, 95% percentile ranking) were ordered by K5 means clustering based on 3D7 (Table S1). Enriched genes compared to previously published RNA-seq analysis (green) (Lopez-Barragan et al., 2011), DNA microarray (identified in ≥ 50% of analyses) (red) (Brancucci et al., 2014; Eksi et al., 2005; Eksi et al., 2012; Mair et al., 2010; Pelle et al., 2015; Silvestrini et al., 2005; Young et al., 2005), and mass spectrometry identification of sexual-stage peptides (blue) (Lindner et al., 2013; Silvestrini et al., 2010; Tao et al., 2014). Genes characteristic of gametocytogenesis and newly identified gametocyte-enriched genes are noted adjacent to the clusters in which they are enriched. B) A subset of 433 genes significantly enriched in F12 (≥ 0.5 log_2_ fold change, 95% percentile ranking) were ordered by K5 means clustering. Genes characteristic of both asexual invasion and gametocytogenesis are labeled. C) Dot-plots representing the median fold change (*pfs16/cam*) of each gene in both 3D7 and F12 transcription and stabilization arrays. Highlighted in blue are gametocyte genes frequently detected (95^th^ percentile) in previously published transcriptomic and proteomic analyses (Table S2 and S3). Example gametocyte gene markers, *pfs16*, *pfg27/25*, *pf14.744*, *adenylate kinase*, and *mdv1* are noted in red.

### Biosynthetic mRNA capture in F12 reveals initiation of gametocyte-specific transcription

Examining the same 808 genes differentially expressed in committed versus non-committed 3D7 parasites, we find that the “transcription” and “stabilization” patterns are largely identical between the *cam*- and *pfs16*-FCU-GFP F12 parasite populations as seen by the overall low fold changes in transcription and stabilization (Fig 5A). For example, Cluster 4 is comprised of genes whose patterns of transcription are invariant between F12^*cam*^ and F12^*pfs16*^, but are highly altered in 3D7^*pfs16*^ gametocytes (Fig. 5A). Of the 808 gametocyte-specific genes identified in 3D7, we find a subset of 431 genes which are significantly (>0.5 log_2_ fold change, 95^th^ percentile) transcribed or stabilized in F12^*pfs16*^ (Fig. 5B). This is interesting, because we know that F12 parasites cannot produce gametocytes and yet we measure the active transcription of these gametocyte-associated genes. These genes include several well-established markers of gametocytogenesis such as *pfpuf2*, *pfs25* and *pfs16* (Fig. 5B, S5A and S5B) (Josling and Llinas 2015) as well the transcription and stabilization of genes involved in merozoite invasion (Fig. 5B, Cluster 4 and 5) {Cowman, 2012 #2078}. Comparison with a list of established *P. falciparum* male and female gametocyte markers (Lasonder et al. 2016) reveals that a large number of dynein genes, NIMA related kinase 4 (*nek4*, PF3D7_0417100/PFD0825c), *Pfgabc2* (PF3D7_1426500/PF14_0244), and *p48/45* 6-cysteine protein (PF3D7_0208900/PF02_0085) are also transcribed in F12^*pfs16*^ (Table S2, Fig. S5A). However, mass spectrometry analysis has shown that these sex-specific gene products do not appear until Stage V of gametocyte development (Silvestrini et al. 2010) (Table S2), and are likely translationally repressed. These data support previous studies demonstrating that male and female sex determination occurs very early on during commitment (Silvestrini et al. 2000; Smith et al. 2000) and that this likely precedes *Pf*AP2-G activity since it is non-functional in F12.

Most notably, we find a weaker gametocyte-specific transcriptional program (involving fewer genes) is initiated in F12 than in 3D7 (Table S2). However, a direct comparison of these genes reveals that they are highly stabilized in 3D7^*pfs16*^ parasites, but not in F12^*pfs16*^ (Fig. 5C and S5B). Therefore, it is likely that in F12, the lack of stabilization of these transcripts, perhaps due to a missing RNA-binding protein may result in an inability to commit to gametocytogenesis. Interestingly, *Pfgexp05* (PF3D7_0936600/PFI1775w), which has been identified as the earliest marker of committed gametocytes (Tiburcio et al. 2015), is actively transcribed during the ring-stage in all strains except 3D7^*pfs16*^ (Fig. S5A), supporting the notion that early gametocyte genes are transcribed independently of the *Pf*AP2-G transcriptional regulator. Taken together, these results suggest that F12 parasites are attempting to commit to gametocytogenesis, but appear to abort prematurely and resume asexual developmental progression possibly due to the lack of stabilization of essential gametocyte-specific transcripts.

### Gametocyte mRNA dynamics support metabolic rewiring and membrane restructuring

Gametocytogenesis is associated with drastic alterations in the morphology, surface-adhesion properties and metabolic functions of the parasite (Liu et al. 2011). To determine if such alterations are predicted by the gametocyte-specific mRNA dynamics we measured, we performed GO-term and KEGG-pathway enrichment analysis on the five gene clusters differentially regulated in *pfs16*-FCU-GFP expressing parasites (Fig. 5A and S6). We find an enrichment of genes involved in membrane structural integrity and transport (Fig. 5A, S6A and S6B; fatty acid metabolism, Clusters 1 and 2; membrane ultrastructure, Clusters 1, 2, and 5; and metabolite transport, Clusters 2 and 5) indicative of the parasite’s preparation to undergo morphological change. Cluster 3 contains genes which are highly transcribed from mtDNA and stabilized post-transcriptionally, including ribosomal subunits and cytochrome c oxidase III (Fig. 5A). Genes encoding additional enzymatic subunits of the mitochondrial electron transport pathway and mitochondrial-associated metabolic pathways are enriched in Cluster 4 (Fig. 5A and S6A) supporting an increase in the mitochondrial metabolic functions of gametocytes (MacRae et al. 2013; Ke et al. 2015). Interestingly, a significant proportion of nuclear encoded tRNAs (24/45) are highly transcribed during this developmental transition (Fig. 5A). Although little is known regarding the role of tRNAs in *Plasmodium* beyond translation, a wide variety of alternative functions have been identified in other eukaryotes (Kirchner and Ignatova 2015). The coordinated upregulation of *Plasmodium* tRNAs warrants further investigation to define their role during gametocytogenesis.

### Gametocyte-specific *cis*-regulatory motif enrichment reveal multi-layered regulation of mRNA dynamics

To predict whether gametocyte mRNA transcription or stabilization is guided by specific DNA/RNA-binding proteins, we performed a DNA sequence motif enrichment analysis on the gametocyte-specific genes in Fig. 5. We found that the DNA motif AGACA was enriched in the 5’ UTR (within 1 kb upstream of the start codon) of a significant proportion (371 of 808 genes, z-score = 191.0, p-value = 1.33e-55) of gametocyte-specific transcribed genes (Fig. S7). This motif has been associated with gametocyte-specific genes in prior bioinformatic analyses of the *Plasmodium* genome and transcriptome (Wu et al. 2008; Young et al. 2008; Russell et al. 2015); however, it has not been associated with any specific transcription factors such as the ApiAP2 DNA-binding proteins (Campbell et al. 2010; Russell et al. 2015). Interestingly, the AP2-G associated motif GTAC was also identified in our search but at a much lower frequency (Fig. S7) (Campbell et al. 2010; Kafsack et al. 2014; Sinha et al. 2014). This suggests that *Pf*AP2-G may regulate a small subset of gametocyte-related genes and additional regulation of gametocyte commitment is required. In the 3’ UTR we detect enrichment of UGUR, which resembles the motif recognized by the *Pf*PUF2 RNA-binding protein (Cui et al. 2002; Miao et al. 2010; Miao et al. 2013) (Fig. S7), indicating that post-transcriptional regulatory mechanisms play a role in gametocytogenesis. The identification of independent regulatory motifs associated with the early gametocyte stage suggests a need for further characterization of the complex gene regulatory network that coordinates mRNA dynamics during this developmental transition, as well as the maturation of the gametocyte.

## DISCUSSION

This study presents a new method to capture and probe transcriptional and post-transcriptional mRNA dynamics in a cell-specific manner at distinct stages of *Plasmodium* development. We have shown that parasites expressing FCU-GFP can readily salvage pyrimidine precursors such as 4-TU, which allows for direct incorporation of this labeled nucleotide into newly transcribed mRNA. Biosynthetic mRNA capture with 4-TU can be genetically tuned to measure either whole populations or stage-specific subpopulations of parasites by altering the promoter driving the expression of FCU-GFP. The separate pools of labeled (transcribed) and unlabeled (stabilized) mRNAs can be isolated from developing *Plasmodium* parasites, and comprehensively measured by DNA microarray analysis (Fig. 1A, 4B and 4C).

In addition to demonstrating the feasibility of this approach in *Plasmodium* parasites, we have applied biosynthetic mRNA capture to examine the dynamic regulation imparted upon transcripts at the developmental transition between the asexual blood-stage and the sexual gametocyte stage. Although previous studies aimed at elucidating the gametocyte transcriptional regulatory program using genome-wide transcriptome analyses have revealed subsets of genes that are differentially abundant during gametocytogenesis (Gissot et al. 2004; Silvestrini et al. 2005; Young et al. 2005; Lopez-Barragan et al. 2011; Pelle et al. 2015), these studies have not measured transcriptional activity or mRNA stability. Rather, they serve to identify genes that have higher transcript abundance during gametocyte development. This study is the first to measure transcription and stabilization of genes necessary at the point of commitment without the need for physical separation of this small subpopulation from the total cell culture. Our results indicate that the mRNA dynamics of commitment to gametocytogenesis consist of a balance of transcription and stabilization in the gametocyte producing line 3D7 (Table S3). Interestingly, we also find that F12 initiates a limited gametocyte-specific transcriptional program (Fig. 5B) which is aborted in each asexual development cycle. Given the weak, but detectable activity of the *pfs16* promoter in F12^*pfs16*^ (Fig. 3C, S2F), these parasites are likely attempting to follow a gametocyte commitment transcriptional program, but due to the lack of a functional AP2-G (Kafsack et al. 2014), they cannot proceed to produce mature gametocytes (Kafsack et al. 2014). Interestingly, biosynthetic capture in F12*^pfs16^* reveals transcription of genes essential to both asexual and sexual development raising the possibility that the mRNA dynamics are more complex than a single factor regulating sexual commitment. Our results also support recent work suggesting a role for commitment events preceding the activity of *Pf*AP2-G (Tiburcio et al. 2015), and suggests a more complex role for AP2-G in gametocytogenesis. However, unlike the gametocyte producing 3D7 parasite line, F12 is not capable of stabilizing transcripts necessary for gametocytogenesis and therefore aborts sexual differentiation. From this study we find that transcription and stabilization of gametocyte specific-genes is an essential determinant of commitment to gametocytogenesis.

We also evaluated the transcriptional dynamics of previously reported gametocyte-specific gene markers and found significant differences between the transcription and stabilization of gametocyte associated genes (Fig. 5A and Table S2). These dynamics allowed us to identify essential metabolic processes and potential DNA regulatory motifs in transcripts associated with gametocyte commitment. While our understanding of gametocyte-specific transcriptional regulation is in its infancy, our data demonstrate that the AP2-G DNA binding motif is associated with several mRNAs transcribed in sexually committed parasites (from *pfs16*-FCU-GFP). However, a majority of gametocyte mRNAs contain an enriched motif (AGACA) for which an associated *trans*-acting factor remains to be identified. Recently, in the murine malaria model, a second ApiAP2 protein *Pb*AP2-G2 has been reported to transcriptionally repress asexual development genes (Yuda et al. 2015). The AP2-G2 motif is distinctly absent from gametocyte-associated genes (Figure 5, Fig. S6) (Campbell et al. 2010; Yuda et al. 2015), suggesting that this repressive role may similarly be conserved in *P. falciparum*. Additionally, our data demonstrate that many markers of gametocytogenesis are stabilized upon commitment, supporting a significant role for translational repression of mRNAs during sexual development. A handful of RNA-binding proteins have been demonstrated to play a role in gametocyte maturation (Cui et al. 2015); however, a specific factor(s) involved in stabilization of transcripts upon commitment to gametocytogenesis remains to be identified.

Although this study has focused on the sexual stage of *Plasmodium*, stage-specific biosynthetic mRNA capture is poised to advance our understanding of parasite transcriptional and post-transcriptional processes at key developmental transitions throughout the entire parasite life cycle. By altering the promoter driving FCU-GFP expression, this method can be adapted to examine mRNA dynamics at any defined point of development using stage-specific promoters from genes known to be upregulated in the ookinete, oocyst, sporozoite, and liver stages. These transcriptional programs can also be determined *in vivo*, since RNA-tagging from insects (Miller et al. 2009), human cell lines (Cleary et al. 2005), and mouse systems (Cleary 2008) have already been established. *In vivo* systems provide further opportunities to explore the relationship between the *Plasmodium* parasite and its diverse host cell types, the mosquito midgut, hepatocytes, and red blood cells. Biosynthetic mRNA capture can also be combined with bioinformatic models to calculate real-time *in vivo* rates of transcription and decay on a whole genome scale (Dolken et al. 2008; Miller et al. 2011; Sun et al. 2012; Neymotin et al. 2014), and may also be used to identify specific responses to environmental perturbations (such as drug-treatment) on short timescales.

We envision that thiol-modified labeling will enhance efforts to define specific or global RNA-protein interactions in the malaria parasite. A recent study identified a large number of RNA-interacting proteins during asexual development; however, the methodology used was not capable of identifying a direct link between these proteins and their target RNAs (Bunnik et al. 2016). Incorporation of 4-TU into the parasite’s mRNA enables photo-activatable UV cross-linking to RBPs that are bound to the thiolated transcripts. This allows for genome-wide RNA-protein interactions to be determined using <u>P</u>hoto<u>a</u>ctivatable-<u>R</u>ibonucleoside-Enhanced Crosslinking (PAR-CL) (Baltz et al. 2012; Castello et al. 2012), or to query the interaction of specific proteins by coupling to an immunoprecipitation step (PAR-CLIP) followed by RNA-sequencing which captures the exact footprint of a specific protein on each transcript (Hafner et al. 2010a; Hafner et al. 2010b). Ultimately, the use of this methodology in the malaria field will advance our understanding of the transcriptional dynamics required for all stages of parasite development, identify regulatory factors involved in gene expression, and enable the identification of new molecular drug targets.

## METHODS

### Transgene construction

The open reading frame of the yeast FCU gene was PCR amplified from *Plasmodium falciparum* vector pCC1 (Maier et al. 2006), and cloned as a translational fusion into the unique BglII and XbaI sites of the pCBM-BSD-684 *5’Pfs16*-gfp vector (Eksi et al. 2008), placing the transcription of the fused gene under the control of a *P. falciparum* gametocyte-specific *pfs16* (PF3D7_0406200/PFD0310w) promoter, yielding the plasmid pCBM-*5’Pfs16*-FCU–GFP. Likewise, FCU was PCR amplified, and cloned into the AvrII and BsiWI sites of the pLN-ENR-GFP (Nkrumah et al. 2006) vector, resulting in pLN-*5’CAM*-FCU-GFP plasmid and placing transcriptional control under the *P. falciparum calmodulin* promotor (PF3D7_1434200/PF14_0323). These plasmids were transformed into, replicated in, and isolated from DH5α *E. coli* for transfection into *P. falciparum*.

### Strains and culture maintenance

*P. falciparum* strains 3D7 (Rovira-Graells et al. 2012) and F12 (Alano et al. 1995) have been described in previous studies and parasite cultures were maintained under standard conditions (Trager and Jensen 1976) at 5% hematocrit of O+ human erythrocytes in RPMI1640 containing hypoxanthine, NaH_2_CO_3_, HEPES, glutamine and 5 g/L AlbuMAX II (Life Technologies).

### Generation of genetically modified parasite lines

Transfection of *P. falciparum* strains was performed as previously described (Fidock and Wellems 1997). Briefly, 5-7% ring-stage parasite cultures were washed three times with 10 times the pellet volume of cytomix. The parasitized RBC pellet was resuspended to 50% hematocrit in cytomix. In preparation for transfection, 100μg of either pCBM-*5’Pfs16*-FCU-GFP or a plasmid was precipitated and resuspended in 100μl cytomix. The plasmid and 250μl of the 50% parasitized RBC suspension were combined and transferred to a 0.2cm electroporation cuvette on ice. Electroporation was carried out using a BioRad GenePulser set at 0.31kV, 960uF. The electroporated cells were immediately transferred to a T-25 flask containing 0.2ml uninfected 50% RBCs and 7ml medium. To select for parasites containing plasmid, medium containing 1.5ug/μl Blasticidin S (Sigma-Aldrich) was added at 48 hours post transfection. Cultures were maintained under constant 1.5ug/μl Blasticidin S pressure, splitting weekly, until viable parasites were observed.

### Verification of transgene expression

Western blot analysis of FCU-GFP protein expression was carried out on mixed stage transgenic *P. falciparum* (10% parasitemia, 5% hematocrit), isolated by saponin (0.01%) lysis. Equal volumes of protein extracts were run on a SDS-10% polyacrylamide gel and transferred to a nitrocellulose membrane. The membrane was blocked with 5% milk. To detect protein, the membrane was exposed to Sheep IgG anti-Cytosine Deaminase antibody primary antibody (Thermo Fisher Scientific) diluted 1:500 in 3% bovine serum albumin and incubated overnight at 4°C followed by a 1 hour incubation with goat-anti sheep-HRP secondary antibody (Thermo Fisher Scientific) diluted 1:1000 in 3% BSA. The membrane was incubated for 1 min in Pierce^®^ ECL-reagent (Thermo Fisher Scientific) and protein detected by exposure to film.

### Assessment of FCU-GFP transgene ability to salvage 4-thioluracil via Northern blot

Function and stage-specific expression of the FCU-GFP transgene was verified in the 3D7^*cam*^ compared to wild-type 3D7 by the addition of various concentrations (0, 10, 20, 40μM) of 4-thiouracil (Sigma Aldrich) to the culture medium from a 200mM stock concentration prepared in DMSO. After parasites were grown in the presence of 4-TU for 12 hours, total RNA was prepared from parasites in 5ml of Trizol. Total RNA was biotinylated using EZ-link Biotin-HPDP (Thermo Fisher Scientific) and 2.5ug run on an Ethidium Bromide 1% agarose gel for 30min at 200V. RNA was transferred to Hybond-N+ nitrocellulose (Amersham) membrane using traditional northern blotting techniques. The RNA was UV-crosslinked to the membrane and probed with streptavidin-HRP (1:1000) (Thermo Fisher Scientific). Biotinylated RNA was detected via incubation with ECL-reagent and exposure to film.

### Thiol-labeling timecourse

*P. falciparum* transgenic parasites were cultured in human erythrocytes at 5% hematocrit in RPMI1640 complete medium as described above. All transgenic parasites were synchronized with three successive (48 hours apart) L-Alanine treatments to remove late stage parasites (Haynes and Moch 2002). To ensure that the resulting population consisted of exclusively asexual parasites, synchronized ring stages were subjected to a 60% Percoll (GE Healthcare) density gradient and any remaining late stages were removed by centrifugation (Wahlgren et al. 1983). These highly synchronous cultures were placed back into culture and allowed to recover for at least 24 hours. At 36 hours post invasion 4-TU was added to the medium at a final concentration of 40μM and parasites were incubated for 12 hours followed by total RNA isolation with TriZol. This 4-TU addition, incubation, and RNA extraction was repeated every 12 hours in 4 successive timepoints throughout intraerythrocytic development.

### Biosynthetic modification and isolation of labeled mRNA throughout development. (*Supplemental Detailed Protocol is available in the Supplementary Materials*)

Briefly, transgenic *P. falciparum* was incubated with 4-TU and total RNA was extracted as described above. All RNA was subjected to the biosynthetic modification and isolation as previously published with a few adjustments (Cleary et al. 2005; Cleary 2008; Zeiner et al. 2008). Specifically, 80μg of total RNA (at a concentration of 0.4μg/μl.) was incubated at room temperature protected from light for 3 hours in the presence of 160 μl of 1mg/ml solution of EZ-link Biotin-HPDP (Thermo Fisher Scientific). Biotinylated total RNA was precipitated and resuspended in DEPC-treated water to a final concentration of 0.5μg/ml. Incorporation of 4-TU and biotinylation was determined by NanoDrop analysis and Northern blot probed with streptavidin-HRP. 4-TU labeled-biotinylated RNA was purified using Dynabeads^®^ MyOne™ Streptavidin C1 magnetic beads (Life Technologies) at a concentration of 2μl/μg of RNA. Beads should be prewashed as per manufacturers protocol to remove any RNases and blocked with 16μg of yeast tRNA (Life Technologies). 4-TU labeled-biotinylated RNA is added to the bead slurry and incubated at room temperature for 20min with rotation. The RNA-bead slurry was placed on a magnetic stand for 1 min, and liquid carefully removed and saved for RNA precipitation. This sample contains RNA that was not thiolated or biotinylated. Then the RNA-bound beads underwent five rounds of stringent washes with buffer consisting of 1M NaCl, 5mM Tris-HCL (pH 7.5), 500μM EDTA in DEPC-treated water. 4-TU labeled-biotinylated RNA was eluted from the beads with 5% 2-mercaptoethanol incubated for 10min, placed back on the magnetic stand and liquid removed and saved for RNA precipitation. The RNA in this fraction contains mRNA transcribed in the presence of 4-TU from FCU-GFP expressing cells. Reverse transcription of RNA and DNA microarray analysis is described in the Supplementary Materians.

Additional experimental and data analysis are provided in Supplementary Information.

## DATA ACCESS

The DNA microarray data have been submitted to Gene Expression Omnibus (https://www.ncbi.nlm.nih.gov/geo/) under study accession number GSE72695.

## AUTHOR CONTRIBUTIONS

Both H.J.P. and M.L. contributed extensively to the work presented in this paper. M.C. provided technical assistance with culture maintenance and strain verification.

## ACKNOWLEDGEMENTS

We would like to thank A.B. Vaidya, Ph.D., Drexel University College of Medicine, Philadelphia, PA, for providing atovaquone, helpful discussions in establishing the experimental methodology and review of the manuscript. We also thank M. Szpara, J. Santos, G. Josling and. S. Lindner for critical reading of this manuscript. We thank the following researchers who provided reagents for this study: K. Williamson (pCBM-BSD-684 *5’Pfs16*-gfp), D. Fidock (pLN-*5’CAM*-FCU-GFP), and P. Alano (*P. falciparum* strain F12). This work was funded through support from the Burroughs Welcome Fund, an NIH Director’s New Innovators award (1DP2OD001315-01), the Center for Quantitative Biology (P50 GM071508), and startup funding from the Pennsylvania State University.

## REFERENCES

Adjalley SH, Johnston GL, Li T, Eastman RT, Ekland EH, Eappen AG, Richman A, Sim BK, Lee MC, Hoffman SL et al. 2011. Quantitative assessment of *Plasmodium falciparum* sexual development reveals potent transmission-blocking activity by methylene blue. PNAS 108: E1214–1223.

Alano P, Roca L, Smith D, Read D, Carter R, Day K. 1995. *Plasmodium falciparum:* parasites defective in early stages of gametocytogenesis. Exp Parasitol 81: 227–235.

Balaji S, Babu MM, Iyer LM, Aravind L. 2005. Discovery of the principal specific transcription factors of *Apicomplexa* and their implication for the evolution of the AP2-integrase DNA binding domains. Nucleic Acids Res 33: 3994–4006.

Baltz AG, Munschauer M, Schwanhausser B, Vasile A, Murakawa Y, Schueler M, Youngs N, Penfold-Brown D, Drew K, Milek M et al. 2012. The mRNA-bound proteome and its global occupancy profile on protein-coding transcripts. Mol Cell 46: 674–690.

Bannister L, Mitchell G. 2003. The ins, outs and roundabouts of malaria. Trends Parasitol 19: 209–213.

Bargieri DY, Thiberge S, Tay CL, Carey AF, Rantz A, Hischen F, Lorthiois A, Straschil U, Singh P, Singh S et al. 2016. Plasmodium Merozoite TRAP Family Protein Is Essential for Vacuole Membrane Disruption and Gamete Egress from Erythrocytes. Cell Host Microbe 20: 618–630.

Baum J, Richard D, Healer J, Rug M, Krnajski Z, Gilberger TW, Green JL, Holder AA, Cowman AF. 2006. A conserved molecular motor drives cell invasion and gliding motility across malaria life cycle stages and other apicomplexan parasites. J Biol Chem 281: 5197–5208.

Bozdech Z, Llinas M, Pulliam BL, Wong ED, Zhu J, DeRisi JL. 2003. The transcriptome of the intraerythrocytic developmental cycle of *Plasmodium falciparum*. PLoS Biol 1: E5.

Brancucci NM, Bertschi NL, Zhu L, Niederwieser I, Chin WH, Wampfler R, Freymond C, Rottmann M, Felger I, Bozdech Z et al. 2014. Heterochromatin protein 1 secures survival and transmission of malaria parasites. Cell Host Microbe 16: 165–176.

Bruce MC, Alano P, Duthie S, Carter R. 1990. Commitment of the malaria parasite Plasmodium falciparum to sexual and asexual development. Parasitol 100 Pt 2: 191–200.

Bruce MC, Carter RN, Nakamura K, Aikawa M, Carter R. 1994. Cellular location and temporal expression of the *Plasmodium falciparum* sexual stage antigen Pfs16. Mol Biochem Parasitol 65: 11–22.

Bunnik EM, Batugedara G, Saraf A, Prudhomme J, Florens L, Le Roch KG. 2016. The mRNA-bound proteome of the human malaria parasite *Plasmodium falciparum*. Genome Biol 17: 147.

Bunnik EM, Chung DW, Hamilton M, Ponts N, Saraf A, Prudhomme J, Florens L, Le Roch KG. 2013. Polysome profiling reveals translational control of gene expression in the human malaria parasite *Plasmodium falciparum*. Genome Biol 14: R128.

Campbell TL, De Silva EK, Olszewski KL, Elemento O, Llinas M. 2010. Identification and genome-wide prediction of DNA binding specificities for the ApiAP2 family of regulators from the malaria parasite. PLoS Pathog 6: e1001165.

Caro F, Ahyong V, Betegon M, DeRisi JL. 2014. Genome-wide regulatory dynamics of translation in the asexual blood stages. Elife 3.

Castello A, Fischer B, Eichelbaum K, Horos R, Beckmann BM, Strein C, Davey NE, Humphreys DT, Preiss T, Steinmetz LM et al. 2012. Insights into RNA biology from an atlas of mammalian mRNA-binding proteins. Cell 149: 1393–1406.

Chene A, Vembar SS, Riviere L, Lopez-Rubio JJ, Claes A, Siegel TN, Sakamoto H, Scheidig-Benatar C, Hernandez-Rivas R, Scherf A. 2012. *Pf*Albas constitute a new eukaryotic DNA/RNA-binding protein family in malaria parasites. Nucleic Acids Res 40: 3066–3077.

Cleary MD. 2008. Cell type-specific analysis of mRNA synthesis and decay in vivo with uracil phosphoribosyltransferase and 4-thiouracil. Methods Enzymol 448: 379–406.

Cleary MD, Meiering CD, Jan E, Guymon R, Boothroyd JC. 2005. Biosynthetic labeling of RNA with uracil phosphoribosyltransferase allows cell-specific microarray analysis of mRNA synthesis and decay. Nat Biotechnol 23: 232–237.

Cowman AF, Berry D, Baum J. 2012. The cellular and molecular basis for malaria parasite invasion of the human red blood cell. J Cell Biol 198: 961–971.

Crabb BS, Cowman AF. 1996. Characterization of promoters and stable transfection by homologous and nonhomologous recombination in *Plasmodium falciparum*. PNAS 93: 7289–7294.

Cui L, Fan Q, Li J. 2002. The malaria parasite *Plasmodium falciparum* encodes members of the Puf RNA-binding protein family with conserved RNA binding activity. Nucleic Acids Res 30: 4607–4617.

Cui L, Lindner S, Miao J. 2015. Translational regulation during stage transitions in malaria parasites. Ann N Y Acad Sci 1342: 1–9.

Dechering KJ, Kaan AM, Mbacham W, Wirth DF, Eling W, Konings RN, Stunnenberg HG. 1999. Isolation and functional characterization of two distinct sexual-stage-specific promoters of the human malaria parasite *Plasmodium falciparum*. Mol Cell Biol 19: 967–978.

Dechering KJ, Thompson J, Dodemont HJ, Eling W, Konings RN. 1997. Developmentally regulated expression of pfs16, a marker for sexual differentiation of the human malaria parasite *Plasmodium falciparum*. Mol Biochem Parasitol 89: 235–244.

Dolken L, Ruzsics Z, Radle B, Friedel CC, Zimmer R, Mages J, Hoffmann R, Dickinson P, Forster T, Ghazal P et al. 2008. High-resolution gene expression profiling for simultaneous kinetic parameter analysis of RNA synthesis and decay. RNA 14: 1959–1972.

Eksi S, Haile Y, Furuya T, Ma L, Su X, Williamson KC. 2005. Identification of a subtelomeric gene family expressed during the asexual-sexual stage transition in *Plasmodium falciparum*. Mol Biochem Parasitol 143: 90–99.

Eksi S, Morahan BJ, Haile Y, Furuya T, Jiang H, Ali O, Xu H, Kiattibutr K, Suri A, Czesny B et al. 2012. *Plasmodium falciparum* gametocyte development 1 (Pfgdv1) and gametocytogenesis early gene identification and commitment to sexual development. PLoS Pathog 8: e1002964.

Eksi S, Suri A, Williamson KC. 2008. Sex- and stage-specific reporter gene expression in *Plasmodium falciparum*. Mol Biochem Parasitol 160: 148–151.

Erbs P, Regulier E, Kintz J, Leroy P, Poitevin Y, Exinger F, Jund R, Mehtali M. 2000. In vivo cancer gene therapy by adenovirus-mediated transfer of a bifunctional yeast cytosine deaminase/uracil phosphoribosyltransferase fusion gene. Cancer Res 60: 3813–3822.

Fidock DA, Wellems TE. 1997. Transformation with human dihydrofolate reductase renders malaria parasites insensitive to WR99210 but does not affect the intrinsic activity of proguanil. PNAS 94: 10931–10936.

Foth BJ, Zhang N, Chaal BK, Sze SK, Preiser PR, Bozdech Z. 2011. Quantitative time-course profiling of parasite and host cell proteins in the human malaria parasite *Plasmodium falciparum*. Mol Cell Proteomics 10: M110 006411.

Foth BJ, Zhang N, Mok S, Preiser PR, Bozdech Z. 2008. Quantitative protein expression profiling reveals extensive post-transcriptional regulation and post-translational modifications in schizont-stage malaria parasites. Genome Biol 9: R177.

Friedel CC, Dolken L. 2009. Metabolic tagging and purification of nascent RNA: implications for transcriptomics. Mol Biosyst 5: 1271–1278.

Gay L, Miller MR, Ventura PB, Devasthali V, Vue Z, Thompson HL, Temple S, Zong H, Cleary MD, Stankunas K et al. 2013. Mouse TU tagging: a chemical/genetic intersectional method for purifying cell type-specific nascent RNA. Genes Dev 27: 98–115.

Gissot M, Refour P, Briquet S, Boschet C, Coupe S, Mazier D, Vaquero C. 2004. Transcriptome of 3D7 and its gametocyte-less derivative F12 *Plasmodium falciparum* clones during erythrocytic development using a gene-specific microarray assigned to gene regulation, cell cycle and transcription factors. Gene 341: 267–277.

Guttery DS, Roques M, Holder AA, Tewari R. 2015. Commit and Transmit: Molecular Players in *Plasmodium* Sexual Development and Zygote Differentiation. Trends in Parasitol doi:10.1016/j.pt.2015.08.002.

Hafner M, Landthaler M, Burger L, Khorshid M, Hausser J, Berninger P, Rothballer A, Ascano M, Jr., Jungkamp AC, Munschauer M et al. 2010a. Transcriptome-wide identification of RNA-binding protein and microRNA target sites by PAR-CLIP. Cell 141: 129–141.

Hafner M, Landthaler M, Burger L, Khorshid M, Hausser J, Berninger P, Rothballer A, Ascano M, Jungkamp AC, Munschauer M et al. 2010b. PAR-CliP--a method to identify transcriptome-wide the binding sites of RNA binding proteins. J Vis Exp doi: 10.3791/2034.

Hall N, Karras M, Raine JD, Carlton JM, Kooij TW, Berriman M, Florens L, Janssen CS, Pain A, Christophides GK et al. 2005. A comprehensive survey of the *Plasmodium* life cycle by genomic, transcriptomic, and proteomic analyses. Science 307: 82–86.

Haynes JD, Moch JK. 2002. Automated synchronization of *Plasmodium falciparum* parasites by culture in a temperature-cycling incubator. Methods Mol Med 72: 489–497.

Hughes KR, Philip N, Starnes GL, Taylor S, Waters AP. 2010. From cradle to grave: RNA biology in malaria parasites. Wiley Interdiscip Rev RNA 1: 287–303.

Hyde JE. 2007. Targeting purine and pyrimidine metabolism in human apicomplexan parasites. Curr Drug Targets 8: 31–47.

Iwanaga S, Kaneko I, Kato T, Yuda M. 2012. Identification of an AP2-family protein that is critical for malaria liver stage development. PloS One 7: e47557.

Josling GA, Llinas M. 2015. Sexual development in *Plasmodium* parasites: knowing when it’s time to commit. Nat Rev Microbiol 13: 573–586.

Kafsack BF, Rovira-Graells N, Clark TG, Bancells C, Crowley VM, Campino SG, Williams AE, Drought LG, Kwiatkowski DP, Baker DA et al. 2014. A transcriptional switch underlies commitment to sexual development in malaria parasites. Nature 507: 248–252.

Ke H, Lewis IA, Morrisey JM, McLean KJ, Ganesan SM, Painter HJ, Mather MW, Jacobs-Lorena M, Llinas M, Vaidya AB. 2015. Genetic investigation of tricarboxylic acid metabolism during the *Plasmodium falciparum* life cycle. Cell Rep 11: 164–174.

Kehrer J, Frischknecht F, Mair GR. 2016. Proteomic Analysis of the *Plasmodium berghei* Gametocyte Egressome and Vesicular bioID of Osmiophilic Body Proteins Identifies Merozoite TRAP-like Protein (MTRAP) as an Essential Factor for Parasite Transmission. Mol Cell Proteomics 15: 2852–2862.

Kenzelmann M, Maertens S, Hergenhahn M, Kueffer S, Hotz-Wagenblatt A, Li L, Wang S, Ittrich C, Lemberger T, Arribas R et al. 2007. Microarray analysis of newly synthesized RNA in cells and animals. PNAS 104: 6164–6169.

Kirchner S, Ignatova Z. 2015. Emerging roles of tRNA in adaptive translation, signalling dynamics and disease. Nat Rev Genet 16: 98-112.

Lasonder E, Rijpma SR, van Schaijk BC, Hoeijmakers WA, Kensche PR, Gresnigt MS, Italiaander A, Vos MW, Woestenenk R, Bousema T et al. 2016. Integrated transcriptomic and proteomic analyses of *P. falciparum* gametocytes: molecular insight into sex-specific processes and translational repression. Nucleic Acids Res 44: 6087–6101.

Le Roch KG, Johnson JR, Florens L, Zhou Y, Santrosyan A, Grainger M, Yan SF, Williamson KC, Holder AA, Carucci DJ et al. 2004. Global analysis of transcript and protein levels across the *Plasmodium falciparum* life cycle. Genome Res 14: 2308–2318.

Le Roch KG, Zhou Y, Blair PL, Grainger M, Moch JK, Haynes JD, De La Vega P, Holder AA, Batalov S, Carucci DJ et al. 2003. Discovery of gene function by expression profiling of the malaria parasite life cycle. Science 301: 1503–1508.

Lelli KM, Slattery M, Mann RS. 2012. Disentangling the many layers of eukaryotic transcriptional regulation. Annu Rev Genet 46: 43–68.

Lindner SE, Mikolajczak SA, Vaughan AM, Moon W, Joyce BR, Sullivan WJ, Jr., Kappe SH. 2013. Perturbations of *Plasmodium* Puf2 expression and RNA-seq of Puf2-deficient sporozoites reveal a critical role in maintaining RNA homeostasis and parasite transmissibility. Cell Microbiol 15: 1266–1283.

Liu Z, Miao J, Cui L. 2011. Gametocytogenesis in malaria parasite: commitment, development and regulation. Future Microbiol 6: 1351–1369.

Llinas M, Bozdech Z, Wong ED, Adai AT, DeRisi JL. 2006. Comparative whole genome transcriptome analysis of three *Plasmodium falciparum* strains. Nucleic Acids Res 34: 1166–1173.

Lopez-Barragan MJ, Lemieux J, Quinones M, Williamson KC, Molina-Cruz A, Cui K, Barillas-Mury C, Zhao K, Su XZ. 2011. Directional gene expression and antisense transcripts in sexual and asexual stages of *Plasmodium falciparum*. BMC Genomics 12: 587.

MacRae JI, Dixon MW, Dearnley MK, Chua HH, Chambers JM, Kenny S, Bottova I, Tilley L, McConville MJ. 2013. Mitochondrial metabolism of sexual and asexual blood stages of the malaria parasite *Plasmodium falciparum*. BMC Biol 11: 67.

Maier AG, Braks JA, Waters AP, Cowman AF. 2006. Negative selection using yeast cytosine deaminase/uracil phosphoribosyl transferase in *Plasmodium falciparum* for targeted gene deletion by double crossover recombination. Mol Biochem Parasitol 150: 118–121.

Mair GR, Lasonder E, Garver LS, Franke-Fayard BM, Carret CK, Wiegant JC, Dirks RW, Dimopoulos G, Janse CJ, Waters AP. 2010. Universal features of post-transcriptional gene regulation are critical for *Plasmodium* zygote development. PLoS Pathog 6: e1000767.

Miao J, Fan Q, Parker D, Li X, Li J, Cui L. 2013. Puf mediates translation repression of transmission-blocking vaccine candidates in malaria parasites. PLoS Pathog 9: e1003268.

Miao J, Li J, Fan Q, Li X, Cui L. 2010. The Puf-family RNA-binding protein *Pf*Puf2 regulates sexual development and sex differentiation in the malaria parasite *Plasmodium falciparum*. J Cell Sci 123: 1039–1049.

Miller C, Schwalb B, Maier K, Schulz D, Dumcke S, Zacher B, Mayer A, Sydow J, Marcinowski L, Dolken L et al. 2011. Dynamic transcriptome analysis measures rates of mRNA synthesis and decay in yeast. Molec Sys Biol 7: 458.

Miller MR, Robinson KJ, Cleary MD, Doe CQ. 2009. TU-tagging: cell type-specific RNA isolation from intact complex tissues. Nat Methods 6: 439–441.

Munchel SE, Shultzaberger RK, Takizawa N, Weis K. 2011. Dynamic profiling of mRNA turnover reveals gene-specific and system-wide regulation of mRNA decay. Mol Biol Cell 22: 2787–2795.

Neymotin B, Athanasiadou R, Gresham D. 2014. Determination of in vivo RNA kinetics using RATE-seq. RNA 20: 1645–1652.

Nkrumah LJ, Muhle RA, Moura PA, Ghosh P, Hatfull GF, Jacobs WR, Jr., Fidock DA. 2006. Efficient site-specific integration in *Plasmodium falciparum* chromosomes mediated by mycobacteriophage Bxb1 integrase. Nat Methods 3: 615–621.

Painter HJ, Campbell TL, Llinas M. 2011. The Apicomplexan AP2 family: Integral factors regulating Plasmodium development. Mol Biochem Parasitol 176: 1–7.

Painter HJ, Morrisey JM, Mather MW, Vaidya AB. 2007. Specific role of mitochondrial electron transport in blood-stage *Plasmodium falciparum*. Nature 446: 88–91.

Pelle KG, Oh K, Buchholz K, Narasimhan V, Joice R, Milner DA, Brancucci NM, Ma S, Voss TS, Ketman K et al. 2015. Transcriptional profiling defines dynamics of parasite tissue sequestration during malaria infection. Genome Med 7: 19.

Pradel G. 2007. Proteins of the malaria parasite sexual stages: expression, function and potential for transmission blocking strategies. Parasitol 134: 1911–1929.

Reddy BP, Shrestha S, Hart KJ, Liang X, Kemirembe K, Cui L, Lindner SE. 2015. A bioinformatic survey of RNA-binding proteins in *Plasmodium*. BMC Genomics 16: 890.

Reyes P, Rathod PK, Sanchez DJ, Mrema JE, Rieckmann KH, Heidrich HG. 1982. Enzymes of purine and pyrimidine metabolism from the human malaria parasite, *Plasmodium falciparum*. Mol Biochem Parasitol 5: 275–290.

Rovira-Graells N, Gupta AP, Planet E, Crowley VM, Mok S, Ribas de Pouplana L, Preiser PR, Bozdech Z, Cortes A. 2012. Transcriptional variation in the malaria parasite *Plasmodium falciparum*. Genome Res 22: 925–938.

Russell K, Emes R, Horrocks P. 2015. Triaging informative cis-regulatory elements for the combinatorial control of temporal gene expression during *Plasmodium falciparum* intraerythrocytic development. Parasit Vectors 8: 81.

Scherf A, Carter R, Petersen C, Alano P, Nelson R, Aikawa M, Mattei D, Pereira da Silva L, Leech J. 1992. Gene inactivation of Pf11-1 of *Plasmodium falciparum* by chromosome breakage and healing: identification of a gametocyte-specific protein with a potential role in gametogenesis. EMBO J 11: 2293–2301.

Silvestrini F, Alano P, Williams JL. 2000. Commitment to the production of male and female gametocytes in the human malaria parasite *Plasmodium falciparum*. Parasitol 121 Pt 5: 465–471.

Silvestrini F, Bozdech Z, Lanfrancotti A, Di Giulio E, Bultrini E, Picci L, Derisi JL, Pizzi E, Alano P. 2005. Genome-wide identification of genes upregulated at the onset of gametocytogenesis in *Plasmodium falciparum*. Mol Biochem Parasitol 143: 100–110.

Silvestrini F, Lasonder E, Olivieri A, Camarda G, van Schaijk B, Sanchez M, Younis Younis S, Sauerwein R, Alano P. 2010. Protein export marks the early phase of gametocytogenesis of the human malaria parasite *Plasmodium falciparum*. Mol Cell Proteomics 9: 1437–1448.

Sinden RE. 1982. Gametocytogenesis of *Plasmodium falciparum* in vitro: an electron microscopic study. Parasitol 84: 111.

Sinha A, Hughes KR, Modrzynska KK, Otto TD, Pfander C, Dickens NJ, Religa AA, Bushell E, Graham AL, Cameron R et al. 2014. A cascade of DNA-binding proteins for sexual commitment and development in *Plasmodium*. Nature 507: 253–257.

Smith TG, Lourenco P, Carter R, Walliker D, Ranford-Cartwright LC. 2000. Commitment to sexual differentiation in the human malaria parasite, *Plasmodium falciparum*. Parasitol 121 (Pt 2): 127–133.

Sun M, Schwalb B, Schulz D, Pirkl N, Etzold S, Lariviere L, Maier KC, Seizl M, Tresch A, Cramer P. 2012. Comparative dynamic transcriptome analysis (cDTA) reveals mutual feedback between mRNA synthesis and degradation. Genome Res 22: 1350–1359.

Tao D, Ubaida-Mohien C, Mathias DK, King JG, Pastrana-Mena R, Tripathi A, Goldowitz I, Graham DR, Moss E, Marti M et al. 2014. Sex-partitioning of the *Plasmodium falciparum* stage V gametocyte proteome provides insight into falciparum-specific cell biology. Mol Cell Proteomics 13: 2705–2724.

Tiburcio M, Dixon MW, Looker O, Younis SY, Tilley L, Alano P. 2015. Specific expression and export of the *Plasmodium falciparum* Gametocyte EXported Protein-5 marks the gametocyte ring stage. Malaria Jl 14: 334.

Trager W, Jensen JB. 1976. Human malaria parasites in continuous culture. Science 193: 673–675.

Vembar SS, Droll D, Scherf A. 2016. Translational regulation in blood stages of the malaria parasite *Plasmodium* spp.: systems-wide studies pave the way. Wiley Interdiscip Rev RNA doi:10.1002/wrna.1365.

Vembar SS, Macpherson CR, Sismeiro O, Coppee JY, Scherf A. 2015. The PfAlba1 RNA-binding protein is an important regulation of translational timing in the *Plasmodium falciparum* blood stages. Genome Biol 16.

Wahlgren M, Berzins K, Perlmann P, Bjorkman A. 1983. Characterization of the humoral immune response in Plasmodium falciparum malaria. I. Estimation of antibodies to *P. falciparum* or human erythrocytes by means of microELISA. Clin Exp Immunol 54: 127–134.

Wu J, Sieglaff DH, Gervin J, Xie XS. 2008. Discovering regulatory motifs in the *Plasmodium* genome using comparative genomics. Bioinformatics (Oxford, England) 24: 1843–1849.

Young JA, Fivelman QL, Blair PL, de la Vega P, Le Roch KG, Zhou Y, Carucci DJ, Baker DA, Winzeler EA. 2005. The *Plasmodium falciparum* sexual development transcriptome: a microarray analysis using ontology-based pattern identification. Mol Biochem Parasitol 143: 67–79.

Young JA, Johnson JR, Benner C, Yan SF, Chen K, Le Roch KG, Zhou Y, Winzeler EA. 2008. In silico discovery of transcription regulatory elements in *Plasmodium falciparum*. BMC genomics 9: 70.

Yuda M, Iwanaga S, Kaneko I, Kato T. 2015. Global transcriptional repression: An initial and essential step for *Plasmodium* sexual development. PNAS 112: 12824–12829.

Zeiner GM, Cleary MD, Fouts AE, Meiring CD, Mocarski ES, Boothroyd JC. 2008. RNA analysis by biosynthetic tagging using 4-thiouracil and uracil phosphoribosyltransferase. Methods Mol Biol 419: 135–146.

